# Volumetric and microstructural regional changes of the hippocampus underlying development of extended delay long-term memory

**DOI:** 10.1101/595827

**Authors:** Anders M Fjell, Markus H. Sneve, Donatas Sederevicius, Øystein Sørensen, Stine K Krogsrud, Athanasia M Mowinckel, Kristine B Walhovd

**Author notes:** Address correspondence to: Anders M Fjell, Dept of Psychology, Pb. 1094 Blindern, 0317 Oslo, Norway.

## Abstract

Episodic memory function improves through childhood and adolescence, in part due to structural maturation of the medial temporal cortex. Although partly different processes support long-term memory over shorter vs. longer intervals, memory is usually assessed after less than an hour. The aim of the present study was to test whether there are unique developmental changes in extended memory, and whether these are related to structural maturation of sub-regions of the hippocampus. 650 children and adolescents from 4.1 to 24.8 years were assessed in total 962 times (mean interval ≈ 1.8 years). Memory was assessed by the California Verbal Learning Test (CVLT) and the Rey Complex Figure Test (CFT). In addition to 30 min recall, an extended delay recall condition was administered ≈ 10 days after encoding. We found unique developmental effects on extended delay memory independently of 30 min recall performance. For visuo-constructive memory, this could be accounted for by visuo-constructive ability levels. Performance was modestly related to anterior and posterior hippocampal volume and mean diffusion. The relationships did not show an anterior-posterior hippocampal axis difference. In conclusion, extended delay memory shows unique development, likely due to changes in encoding depth or efficacy, or improvements of long-term consolidation processes.

**Highlights:**

- Unique developmental effects on episodic memories over days rather than minutes
- Development of visuoconstructive recall explainable by visuoconstructive abilitity
- Development of verbal recall cannot be explained by verbal ability
- Modest relationships between memory and hippocampal structural features

## 1.0 Introduction

Episodic memory function improves through childhood and adolescence [1], likely partly due to structural maturation of critical brain regions such as the medial temporal and the prefrontal cortex [2, 3]. Of special importance for episodic memory is the hippocampus. Hippocampus is necessary for encoding of new information [4], and engaged in retrieval of vivid episodic memories [5]. Hippocampus also plays a unique role in consolidation, maintenance and transformation of episodic memories [6]. Accordingly, studies have shown stronger relationships between hippocampal volume and memory over days and weeks compared to hours or less in development [2], adulthood and aging [7]. Studies of hippocampal-neocortical connectivity also point to stronger involvement of hippocampus for encoding of durable episodic memories [8].

The aim of the present study was to test whether there are unique developmental improvements in extended memory performance, and whether structural development of sub-regions of the hippocampus is related to this. Different hippocampal sub-regions are involved in processing of episodic memory information, with partly distinguishable contributions [6, 9, 10]. Much focus has been on a long axis anterior-posterior (AP) gradient [11–13], where functional studies have suggested encoding to be supported relatively more by the anterior part of hippocampus (aHC) and retrieval relatively more by the posterior (pHC) [11, 13–16]. This may also be relevant for development of memory, since partly different age-trajectories along the AP-axis have been found [3, 17–21]. Relationships between memory performance and the volume of anterior-posterior sub-regions have also been reported to differ between children and adults. One study found that the volume of the posterior hippocampus was related to memory performance in children while anterior volume [3] and activity [22] was related in adults, the latter in accordance with results of a large adult study [23]. The literature is not consistent, however, since other studies have found memory-volume correlations in the anterior hippocampus in children [24] and the posterior in young adults [25].

Volumetric changes may reflect various cellular processes within the hippocampus, such as neurogenesis [26], non-neuronal cell changes [27], cell death and synaptic changes [28, 29], pruning [30], myelination [31] and vascularization [32]. Several of these processes may alter water diffusion in the tissue, which can be measured by diffusion tensor imaging (DTI). Higher mean diffusivity (MD) in the hippocampus is related to aging [33–35], increases over time in older adults [36] and correlates more strongly negatively with memory function than does hippocampal volume [34, 37–39]. Both macro- and microstructural properties of the hippocampus may contribute to explain developmental changes in episodic memory function. In our previous life-span study, using a sample overlapping the present [40], we found relationships between both hippocampal macro- and microstructure and episodic memory scores over 30 min intervals, but these were dependent on the common influence from age.

In the present study, our first aim was to test whether extended delay memory, i.e. episodic memories tested on average 10 days after encoding, showed unique developmental trajectories that could not be accounted for by more immediate long-term memory, i.e. memory as conventionally tested 30 minutes after encoding. In a previous cross-sectional study of 8 to 19 year olds, we found no developmental effects on visuo-constructive 1 week recall when 30 min recall performance was accounted for [2]. Performance on the extended delay memory test still correlated with total hippocampal volume. In the present study, we use a larger sample (n = 650 vs. n = 107), including longitudinal observations (312 longitudinal examinations) and a wider age-range (4.1 to 24.8 vs. 8-19 years). This allowed us to assess developmental changes with higher sensitivity and superior statistical power. In addition, we tested both visuo-constructive and verbal recall.

Our second aim was to relate development of extended delay memory to structural maturation of sub-regions of the hippocampus. Hippocampus was divided in an anterior (aHC) and a posterior (pHC) part according to established procedures [11]. Macro-structural maturation was measured as regional volume and micro-structural maturation as regional mean diffusion (MD). We hypothesized that unique developmental trajectories would be seen for extended delay memory, and that memory performance would be positively related to regional hippocampal volume and negatively to regional hippocampal MD. Whether aHC or pHC would be more strongly related to memory is an open question, as previous results have not been consistent. Since connectivity studies have revealed distinct connections between the hippocampal regions and the neocortex, and there are few direct connections between aHC and pHC [11], it is conceivable that they may play different roles in the hippocampal-neocortical replay necessary for memory over extended time intervals.

## 2.0 Materials and methods

### Sample

Participants were drawn from studies coordinated by the Research Group for Lifespan Changes in Brain and Cognition (LCBC www.oslobrains.no) [41], approved by a Norwegian Regional Committee for Medical and Health Research Ethics. Written informed consent was obtained from all participants older than 12 years of age and from a parent/guardian of volunteers under 16 years of age. Oral informed consent was obtained from participants under 12 years of age. The full sample consisted of 650 healthy participants, 4.1 - 24.8 years of age (mean examination age, 10.4 years years, 1^st^ quartile = 6.7 years, 3^rd^ quartile = 12.9) with a total of 914 MRI examinations and up to 832 memory tests sessions (832 observations for CVLT 30 min recall; 770 for CVLT extended delay recall; 666 for CFT 30 min; 602 for CFT extended delay). Participants were followed for up to 4 time points with MRI, for a maximum period of 8.9 years since baseline (mean interval between visits = 1.8 years, mean total follow up time since baseline for the longitudinal examinations = 1.9 years). Adult participants (> 20 years) were screened using a standardized health interview prior to inclusion in the study (see [40]). Participants with a history of self- or parent-reported neurological or psychiatric conditions, including clinically significant stroke, serious head injury, untreated hypertension, diabetes, and use of psychoactive drugs within the last two years, were excluded. Further, participants reporting worries concerning their cognitive status, including memory function, were excluded.

### Memory testing

The California Verbal learning Test (CVLT) [42] was used to assess verbal episodic memory and the Rey-Osterrieth Complex Figure (CFT) test [43] was used to assess visuo-constructive episodic memory. To allow comparison between verbal and visual memory, scores on the 30 minutes recall conditions were used for both tests.

The learning part of CVLT consists of oral presentation of 16 words in four semantical categories, and the whole list is presented five times with a free recall trial after each presentation. After five presentations, the free recall trials are repeated 5 and 30 minutes later. In addition, an additional extended delay recall condition was administered after a mean delay of 9.9 days (1^st^ quartile = 7 days, 3^rd^ quartile = 10 days).

For visual memory, CFT measures visuo-constructive memory using a novel, complex design which participants are asked to copy and then reproduce from memory after 30 min. The participants were presented with a picture of a geometrical figure on an A4 sheet of paper and were asked to draw the figure as similar as possible. After approximately 30 min, during which time the participants completed other tasks with mainly verbal material, they were asked to draw the figure again without the original picture in front of them. The scoring system divides the figure into 18 subunits and awards 2 points for each correct and correctly placed unit, 1 point for an inaccurately drawn or incorrectly placed unit, and a 1/2 point for a unit that is recognizable but both inaccurate and inaccurately placed in the drawing. This results in a maximum score of 36 points for each drawing. As for CVLT, an extended delay recall condition was administered after a mean delay of 10.2 days (1^st^ quartile = 7 days, 3^rd^ quartile = 10 days).

### MRI acquisition and cross-scanner validation

858 scans were obtained from 1.5 Tesla Avanto (12 channel head coil) and 56 from 3T Skyra (20 channel head coil). The following sequences were used:

Avanto T1-weighted: 2 repeated 3D T1-weighted magnetization prepared rapid gradient echo (MPRAGE): TR/TE/TI = 2400 ms/ 3.61 ms/ 1000 ms, FA = 8°, acquisition matrix 192 × 192, FOV = 240×240 mm, 160 sagittal slices with voxel sizes 1.25 × 1.25 × 1.2 mm. For most children 4-9 years old, iPAT was used, acquiring multiple T1 scans within a short scan time, enabling us to discard scans with residual movement and average the scans with sufficient quality.

Avanto DTI: 32 directions, TR = 8200 ms, TE = 81 ms, b-value = 700 s/mm2, voxel size = 2.0 × 2.0 × 2.0 mm, field of view = 128, matrix size = 128 × 128 × 64, number of b0 images = 5, GRAPPA acceleration factor = 2.

Skyra T1- weighted: 176 sagittal oriented slices were obtained using a turbo field echo pulse sequence (TR = 2300 ms, TE = 2.98 ms, flip angle = 8°, voxel size = 1 × 1 × 1 mm, FOV = 256 × 256 mm). For the youngest children, integrated parallel acquisition techniques (iPAT) was used, acquiring multiple T1 scans within a short scan time, enabling us to discard scans with residual movement and average the scans with sufficient quality.

Skyra DTI: A single-shot twice-refocused spin-echo echo planar imaging (EPI) with 64 directions: TR = 9300 ms, TE = 87 ms, b-value = 1000 s/mm2, voxel size = 2.0 × 2.0 × 2.0 mm, slice spacing = 2.6 mm, FOV = 256, matrix size = 128 × 130 × 70, 1 non-diffusion-weighted (b = 0) image. A b0-weighted image was acquired with the reverse phase encoding.

Since different scanners will yield different volumetric and MD values, 180 participants evenly distributed across a wide age range were scanned on the 1.5T (Avanto) scanner and the 3T (Skyra) scanner on the same day, to allow us to directly assess the influence of scanner. These data are previously published [40], showing that the different scanners yielded highly significant differences in absolute MD and volume. The correlations between scanners were good, however, r = .85 (anterior) and .88 (posterior) for volume and .71 (anterior) and .73 (posterior) for MD. The high rank-order coherence between scanners suggested that inclusion of scanner as a covariate in the analyses efficiently would remove most of the variance between participants due to different scanners.

### MRI preprocessing - morphometry

T1-weighted scans were run through FreeSurfer 6.0 (https://surfer.nmr.mgh.harvard.edu/). FreeSurfer is an almost fully automated processing tool [44–47], and manual editing was not performed to avoid introducing errors. For the children, the issue of movement is especially important, as it could potentially induce bias in the [48]. Rather, all scans were manually rated for movement on a 1-4 scale, and only scans with ratings 1 and 2 (no visible or only very minor possible signs of movement) were included in the analyses, reducing the risk of movement affecting the results. Also, all reconstructed surfaces were inspected, and discarded if they did not pass internal quality control. The hippocampus was initially segmented as part of the FreeSurfer subcortical stream [45] before being divided in aHC and pHC (see below). 90 scans were discarded due to low quality due mainly to excessive motion, technical issues during acquisition, incomplete protocols (e.g. lacking DTI data) or reconstruction or segmentation issues, reducing the number of scans in the analyses to 824 for both volume and mean diffusion (see below).

### MRI preprocessing – DTI

DTI scans were processed with FMRIB’s Diffusion Toolbox (fsl.fmrib.ox.ac.uk/fsl/fslwiki) [49, 50]. B0 images were also collected with reversed phase-encode blips, resulting in pairs of images with distortions going in opposite directions. From these pairs we estimated the susceptibility-induced off-resonance field using a method similar to what is described in (Andersson et al., 2003) as implemented in FSL (Smith et al., 2004). We then applied the estimate of the susceptibility induced off-resonance field with the eddy tool (Andersson and Sotiropoulos, 2016), which was also used to correct eddy-current induced distortions and subject head movement, align all images to the first image in the series and rotate the bvecs in accordance with the image alignments performed in the previous steps (Jenkinson et al., 2002; Leemans and Jones, 2009).

### Hippocampal anterior-posterior segmentation

Moving anteriorly through the coronal planes of an MNI-resampled human brain, y = −21 corresponds to the appearance of the uncus of the parahippocampal gyrus. In line with recent recommendations for long-axis segmentation of the hippocampus in human neuroimaging (Poppenk et al., 2013), we labeled hippocampal voxels at or anterior to this landmark as anterior HC while voxels posterior to the uncal apex were labeled as posterior HC. Specifically, for each participant, all diffusion voxels for which more than 50% of the underlying anatomical voxels were labeled as hippocampus by FreeSurfer [45] were considered representations of the hippocampus. While keeping the data in native subject space, we next established hippocampal voxels’ locations relative to MNI y = −21 by calculating the inverse of the MNI-transformation parameters for a given subject’s brain and projecting the back-transformed coronal plane corresponding to MNI y = −21 to diffusion native space. All reported diffusion measures thus represent averages from hippocampal sub-regions established in native space. An illustration of this segmentation is shown in Figure 1.

**Figure 1.**
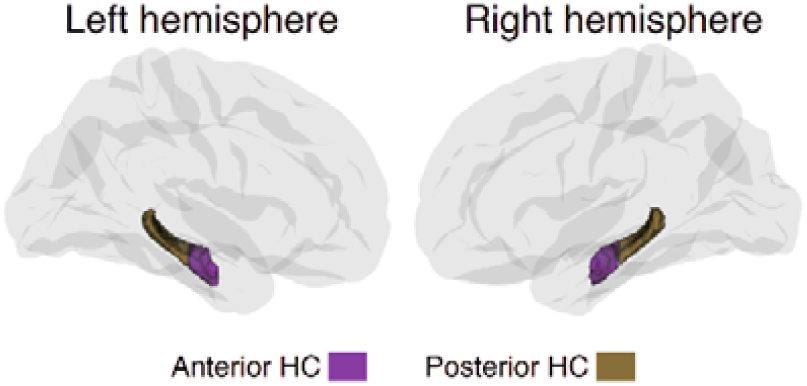
Hippocampal sub-regions. Hippocampus was segmented in an anterior and a posterior part according to established procedures.

### Statistical analyses

Analyses were run in R (https://www.r-project.org) using Rstudio (www.rstudio.com) IDE. Generalized Additive Mixed Models (GAMM) using the package “mgcv” [51] were used to derive age-functions. Different models were run with the different memory recall scores in turn as dependent variables. We included a smooth term for age, random effect for subject, and sex as covariate of no interest. Subject time-point was included as an additional covariate in all memory analyses to control for practice effects of the memory scores. To test for unique developmental trajectories of extended delay memory, extended delay memory was used as dependent variable and 30 min recall as an additional covariate. If the relationship between extended delay memory and age was still significant, this was taken as evidence of a unique developmental effect on extended delay memory. Then each hippocampal sub-region and modality was used as dependent variables in separate analyses. Scanner was used as an additional covariate of no interest in all analyses of hippocampal sub-regions. To assess the relationship between hippocampal sub-regions and memory function, age and hippocampal sub-regions were included in the same models, with the same covariates as above.

In all the models, the smoothness of the age-curve is estimated as part of the model fit, and the resulting effective degrees of freedom (edf) was taken as a measure of deviation from linearity. The p-values associated with the smooth terms are only approximate, as they are based on the assumption that a penalized fit is equal to an unpenalized fit with the same edf, and do not take into account uncertainty associated with the smoothing parameter estimation. The major advantage of GAMM in the present setting is that relationships of any degree of complexity can be modelled without specification of the basic shape of the relationship, and GAMM is thus especially well-suited to map life-span trajectories of neurocognitive variables which can be assumed to be non-linear and where the basic form of the curve is not known (Fjell et al., 2010).

Since not all information was available for all participants, the number of observations that were used in each analysis is presented.

## 3.0 Results

### Age-relationships

Developmental memory trajectories are presented in Figure 2. 30 min recall scores showed the expected sharp development through childhood for both CVLT (F = 158.6, edf = 5.0, p < 2e^−16^, n = 832) and CFT (F = 56.6, edf = 43.9, p < 2e^−16^, n = 666). However, the trajectories still differed substantially. While CVLT score increased throughout the age-span, CFT peaked in mid adolescence. Next, the same analyses were run for the extended delay memory variables. Both CVLT (F = 156.9, edf = 4.9, p < 2e^−16^, n = 752) and CFT (F = 57.9, edf = 3.4, p = 2e^−16^) developed rapidly through childhood and adolescence. For CVLT, however, the developmental trajectory for the extended delay memory performance showed an earlier and steeper peak than the 30 min performance, suggesting developmental differences. Thus, the GAMMs were re-run with 30 min performance as an additional covariate in the age-analyses of the extended delay memory scores. These analyses showed clear effects of age on extended delay recall, independently of 30 min recall, both for CVLT (F = 64.7, edf = 5.1, p < 2e^−16^) and CFT (F = 31.6, edf = 1, p = 2.9e^−08^). For CFT, the residual age-trajectory of extended delay memory was linear and positive when 30 min performance was accounted for. For CVLT, there was no additional developmental improvements over and above 30 min recall until about 10 years, from which sharp positive development was seen until the last part of adolescence.

**Figure 2.**
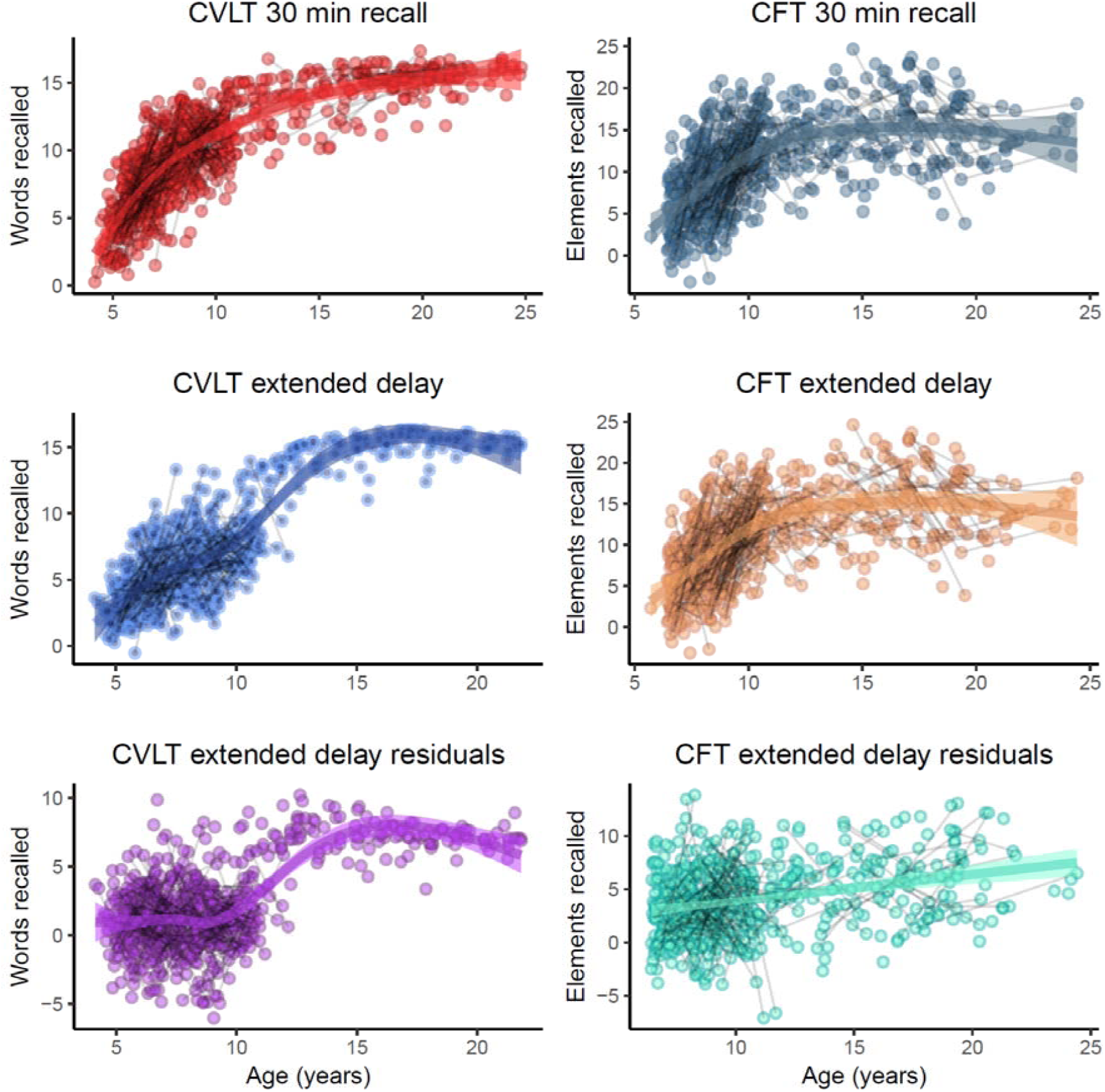
Developmental trajectories for memory. Development of verbal (left column) and visuo-constructive (right column) recall performance. The plots in the bottom row show how performance in the extended delay 10 days recall condition improves when performance on the 30 min condition is accounted for.

### Relationships to hippocampal volume and macrostructure

Developmental trajectories for volume and MD for hippocampal sub-regions are shown in Figure 3. These are included as background information, since the current data have previously been used in a separate publication on a larger sample [40]. As expected, hippocampal anterior (F = 38.5, edf = 3.7, p < 2e^−16^, n = 824 for all analyses) and posterior (F = 16.4, edf = 3.8, p = 4.8e^−12^) volume increased early in development, but either peaked towards the end of adolescence (anterior) or flattened out showing only modest changes from early teen years (posterior). For MD, the relationships differed substantially in the anterior vs. posterior regions. Anterior MD was reduced in childhood, and showed little further change during adolescence (F = 13.9, edf = 2.9, p = 9.3e^−09^). In contrast, posterior MD did not show any significant developmental effect (F = 2.7, edf = 1, p = .10). These results demonstrate the volume and MD show different developmental trajectories, and that the anterior and posterior region of the hippocampus differ in development.

**Figure 3.**
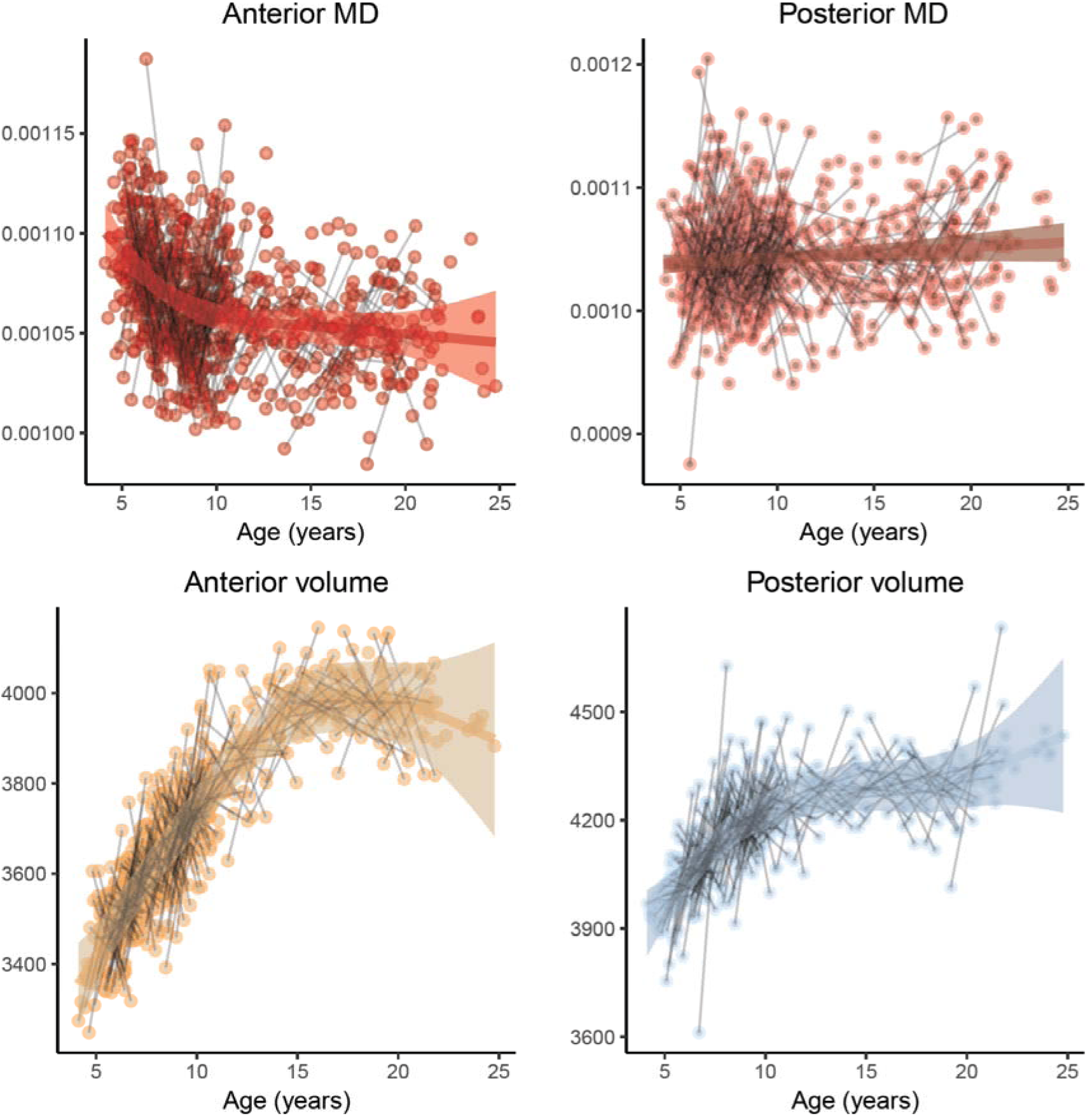
Developmental trajectories for hippocampus. Structural maturation of hippocampal sub-regions. Top row shows microstructure (mean diffusion), bottom row shows volume.

Finally, we tested the relationship between memory performance and hippocampal sub-regional volume, controlling for age, sex, subject time point and scanner (number of observations for each analysis: CVLT 30 min n = 715; CVLT extended delay n = 663; CFT 30 min n = 615; CFT extended delay n = 555). The results are presented in Table 1 and Figure 4 and illustrated with scatterplots in Figure 5. Four relationships were found at an uncorrected α-level of .05. The only relationship that survived full Bonferroni corrections was between CVLT extended delay memory performance and anterior hippocampal MD, with lower MD being associated with higher memory score. This relationship was specific to extended delay recall, as it was not found for the 30 min condition, and still was significant when adding 30 min recall as an additional covariate (p = .014). We ran an additional GAMM adding an age x aHC MD interaction term, which was not significant (p = .61).

**Table 1.** Relationship between hippocampal sub-regions and memory **Bold** indicate p < .05, Bonferroni corrected for 16 comparisons ***Bold italics*** indicate p <.05 uncorrected

| CVLT |  |  |  |  |  |  |
| --- | --- | --- | --- | --- | --- | --- |
| 30 min recall |  |  |  | Extended delay recall |  |  |
|  | edf | F | p < | edf | F | p < |
| Ant vol | 1.0 | 3.1 | .08 | 1.0 | 5.2 | **.023** |
| Post vol | 1.0 | 0.2 | .69 | 1.0 | 0.3 | .60 |
| Ant MD | 1.0 | 0.9 | .37 | 1.0 | 7.7 | **.006** |
| Post MD | 1.0 | 0.4 | .54 | 1.0 | 1.9 | .17 |
| CFT |  |  |  |  |  |  |
| 30 min recall |  |  |  | Extended delay recall |  |  |
|  | edf | F | p < | edf | F | p < |
| Ant vol | 1.0 | 2.5 | .11 | 1.0 | 4.0 | .11 |
| Post vol | 1.0 | 1.5 | .22 | 1.0 | 6.6 | **.01** |
| Ant MD | 2.4 | 5.1 | **.01** | 1.0 | 0.3 | .60 |
| Post MD | 1.0 | 0.1 | .77 | 1.0 | 2.3 | .13 |

**Figure 4.**
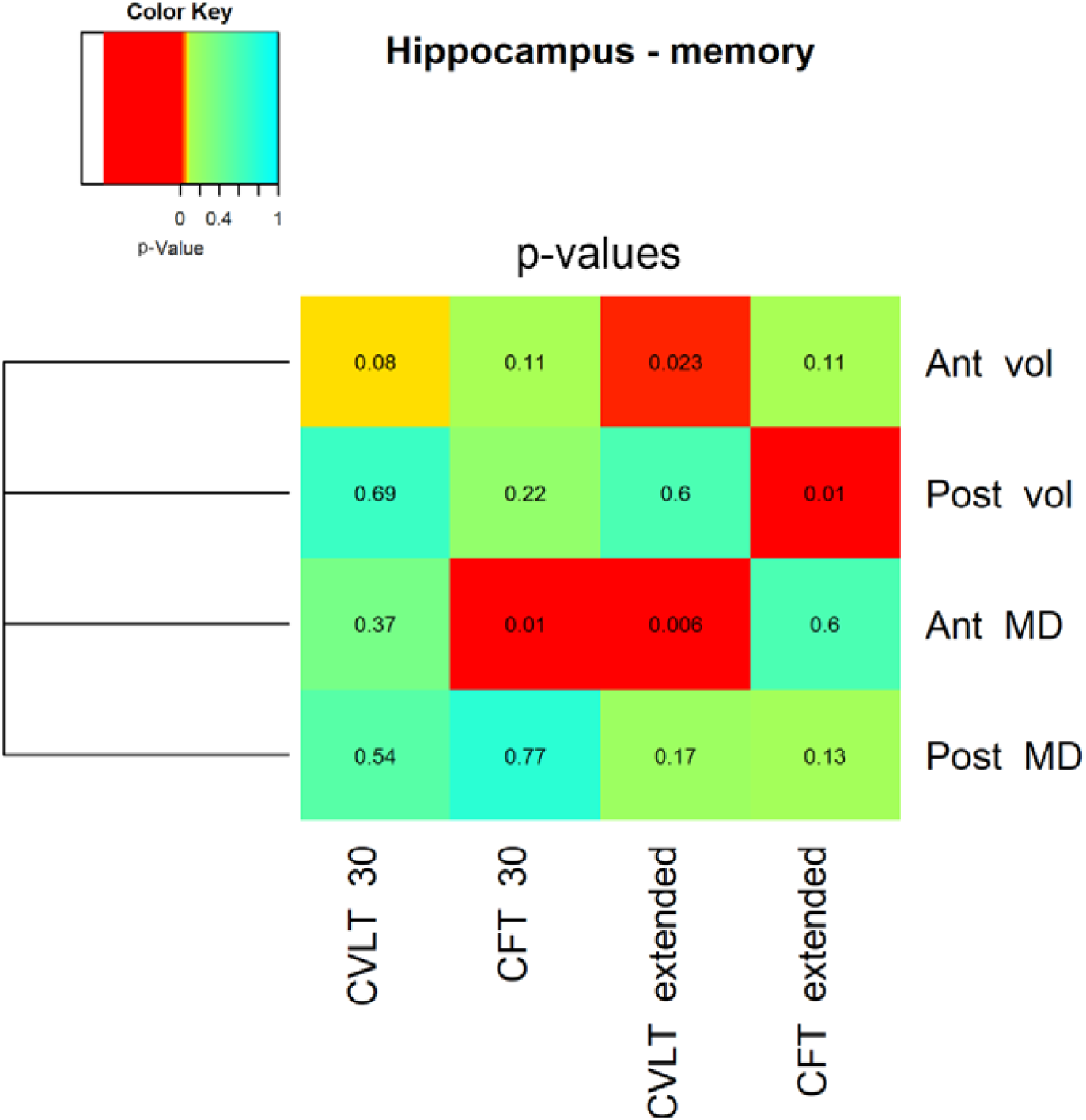
Memory and hippocampal subfields p-values. The heat plot illustrates the statistical significance (uncorrected p-value) of the relationship between hippocampal structure (volume and mean diffusion) and memory performance. Age was included as a covariate in the analyses.

**Figure 5.**
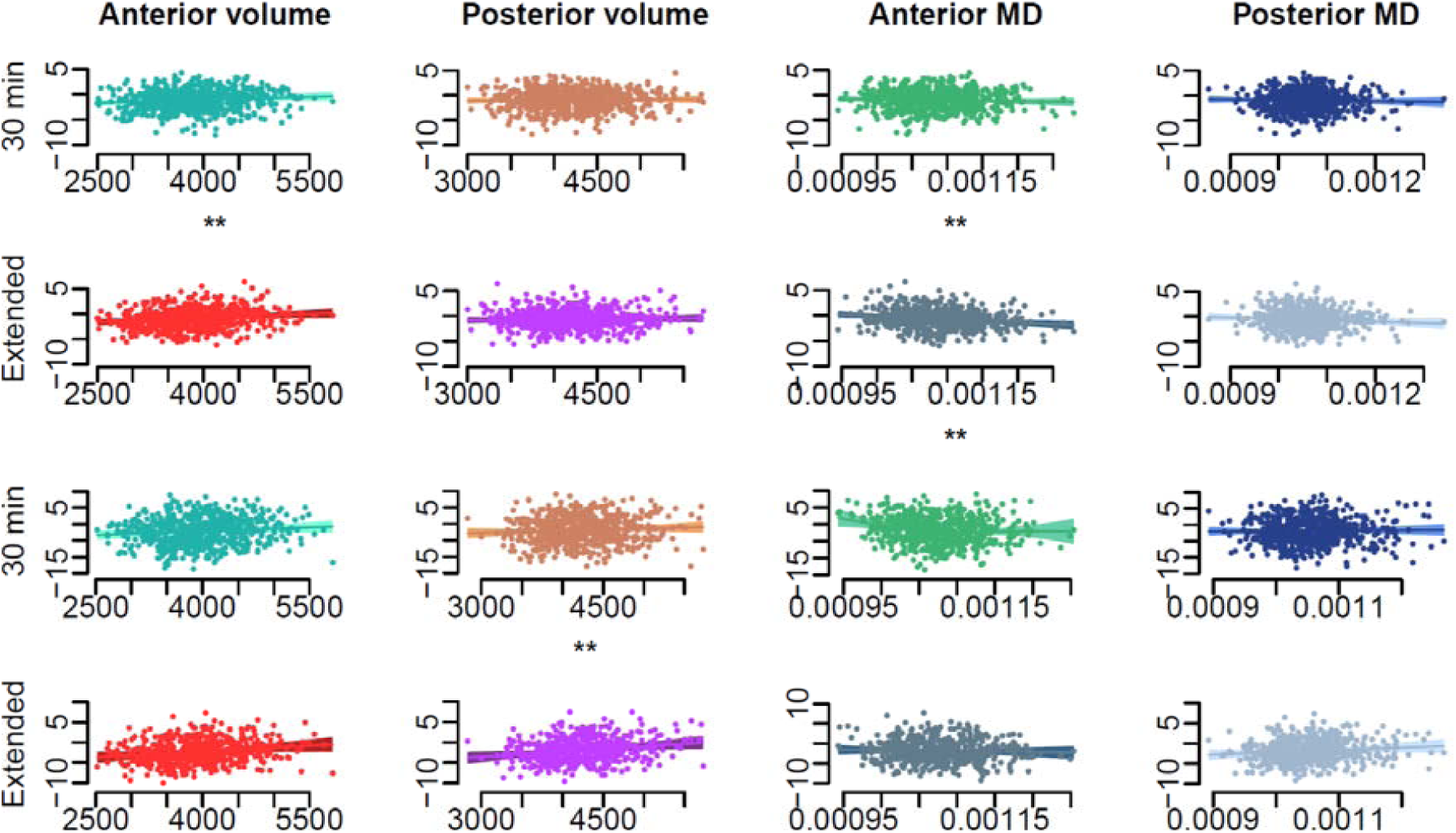
Memory and hippocampal subfields relationships. Scatterplots illustrating the relationship between hippocampus structure and memory performance. The two top rows show verbal recall performance. The two bottom rows show visuo-constructive performance. ** p < .05

### Influence of development of verbal and visuo-constructive abilities

We were interested in testing whether the developmental effects on the extended delay memory scores were related to development of general verbal (for CVLT) and visuo-constructive (for CFT) abilities. Thus, we re-ran the GAMM functions with CVLT extended delay score as dependent and age as a smooth predictor, with 30 min recall, sex, retention interval and time point as covariates. In addition, we added similarities and vocabulary raw scores from WASI. The relationship between age and extended delay CVLT score was still highly significant (F = 27.3, edf = 6.8, p < 2e^−16^, n = 737), while neither of the WASI variables contributed (both p’s > .15). Since the WASI variables are expected to be highly correlated, we re-ran the analysis with vocabulary and similarities in separate models. Now vocabulary contributed significantly and positively, although modestly, to extended delay recall score (vocabulary: t = 2.2, p = .03, n = 741; similarities: t = 1.8, p = .076, n = 737). Formally testing whether verbal abilities significantly affected the developmental trajectory of CVLT extended delay recall, we calculated the residual age-function with and without the WASI variables as covariates. We then tested whether the derivatives of the models differed at any age. This was not the case, which implies that verbal ability levels do not significantly affect the age-trajectory of extended delay verbal memory (see Figure 6).

**Figure 6.**
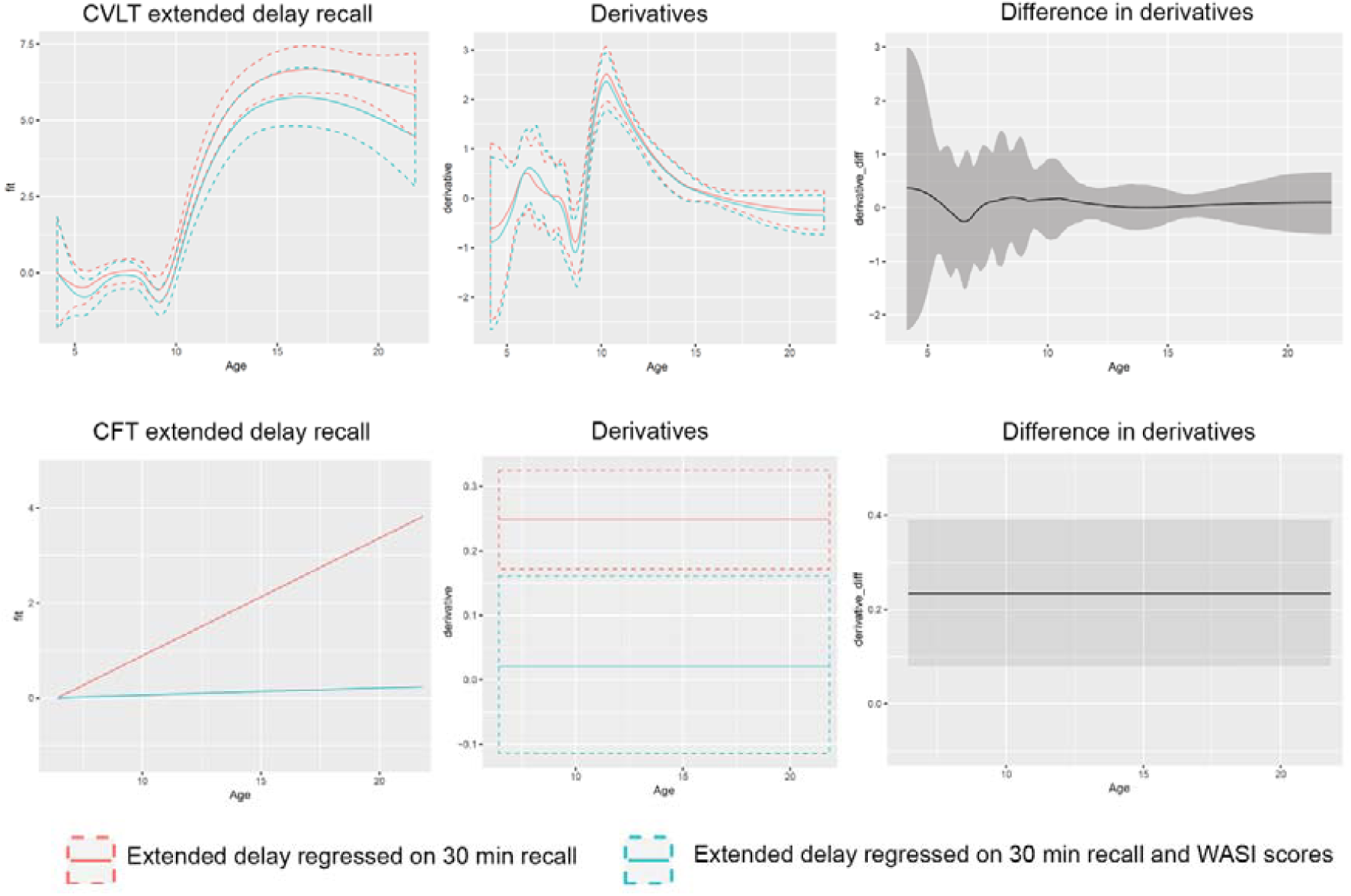
Effects of verbal and visuo-constructive abilities. We tested how verbal and visuo-constructive ability level affected the age-relationship of extended delay memory recall. Top row shows the result for verbal recall. Bottom row shows the results for visuo-constructive recall. In the first column, the age-trajectory without ability level controlled for is shown in red, and the age-trajectory with ability level controlled for in blue-cyan. In the middle row, the respective derivatives of the age-curves are shown. In the right column, the differences in derivatives between the red and the blue-cyan curves are plotted. When the confidence interval of these differences does not include zero, this means that the effect of ability level on the age-trajectory is significant. This is the case for visuo-constructive extended delay recall (bottom right corner), but not extended delay verbal recall (top right corner).

The same type of analysis was run with CFT extended delay recall as dependent and the WASI measures of block design and matrix reasoning as covariates. Inclusion of these rendered the effect of age not significant (F = 0.06, edf = 1, p = .81, n = 591). When tested in separate models, both block design (t = 4.9, p = p < 1.25e^−6^, n = 591) and matrix reasoning (t = 3.2, p = .001, n = 599) were significantly and positively related to extended delay memory performance. As for CVLT, we performed a formal test of the effects of the WASI tests on the developmental trajectory of CFT extended delay recall. Testing the derivatives of the age-function with vs. without the WASI tests as covariates revealed a significant influence on the age-trajectory (see Figure 6).

## 4.0 Discussion

Three main conclusions can be drawn from the present study. First, there are distinct developmental trajectories for extended delay episodic memory that cannot be explained by long-term memory performance over shorter time intervals. Second, memory performance over extended intervals was modestly related to structural features of the hippocampus, with only anterior microstructure contributing to delayed memory score using strict corrections. Finally, visuo-constructive ability level significantly affected the developmental trajectory of extended delay visual recall, explaining the major part of the age effect, while the developmental trajectory of extended delay verbal recall was not affected by verbal ability level. The implications of the findings are discussed below.

### Development of long-term memory

Although a primary function of episodic memory is to keep information in an accessible form over prolonged intervals, long term memory is usually tested after an hour or less. This is based on a premise that the processes responsible for long-term memory can be evaluated after short time intervals. However, here is good evidence from neuropsychological [52] and molecular studies [53] that episodic information is stored in long-term memory through an initial process of rapid consolidation followed by a slower consolidation phase, and that these consolidation phases depend on fundamentally different processes in the brain. Hippocampal-neocortical replay is ongoing for extended time after encoding [6], and both hippocampus and its structural and functional connections show substantial developmental changes through childhood [16, 54–57]. Thus, it would not be surprising if these extended consolidation processes are affected by maturational events in childhood and adolescence. Developmental effects on the processes responsible for stabilization, maintenance and transformations of episodic memories over longer time intervals would be expected to result in specific developmental effects on delayed memory functions that cannot be accounted for by long-term memory performance over shorter time. This was exactly what was found in the present study, for two very different recall tasks. There were unique developmental effects on both extended delay verbal and visuo-constructive memory, over and above the 30 min recall conditions. In adults, the phenomenon of accelerated long-term forgetting (ALF) has been used to refer to abnormal forgetting over hours to weeks despite normal encoding or initial consolidation [58, 59]. This suggests that episodic memory over longer time intervals is supported by partly separate brain processes than memory over 30 minutes, which is in line with the unique developmental effects observed for the extended delay recall conditions in the present study. Similarly, although the evidence is partly mixed, there seems to be increased forgetting over long time intervals also in the other end of the lifespan [58].

Importantly, however, it is a fallacy to conclude from the present results alone that consolidation processes are selectively changing through development. Brain activation studies have shown that hippocampal-neocortical connectivity during encoding can explain differences between memory performance over hours compared to weeks [8]. In one such study it was found that a critical level of activation of the hippocampus during encoding was necessary for source memory performance for both hours and weeks, while strong levels of connectivity between hippocampus and perceptual and self-referential default mode networks during encoding were necessary for establishment of durable source memories [8]. This is relevant for the interpretation of the present results, because these effects were observed at the stage of encoding, in principle independently of extended consolidation processes. It is likely that the increase in hippocampal-cortical connectivity seen during encoding also affected consolidation processes over extended intervals, but this is a speculation. Similarly, we do not know to what degree the unique developmental trajectories for recall performance over extended time intervals is due to differences in encoding or consolidation, or, most likely, a combination of them.

The present results also showed that although verbal ability level as measured by the WASI subtest vocabulary was modestly related to verbal recall performance over extended delay, the age-trajectory was minimally affected by verbal ability level. In contrast, development of visuo-constructive extended delay recall performance was highly influenced by scores on the matrix reasoning and block design tests, especially the latter. Controlling for performance on these two tests completely removed the developmental effect on extended delay memory. This is in line with previous observations that the copy score on the CFT improves significantly in development and is highly related to short- and long-term recall [2]. The strong effects of visuo-constructive abilities on extended delay recall performance may be an instance of statistical collinearity, and the developmental effects on extended delay recall may be real but impossible to disentangle from developmental effects in other cognitive domains. However, the effects may also be caused by higher visuo-constructive abilities leading to more efficient or elaborative encoding strategies due to better understanding and conceptualization of the material, again causing superior organization and hence improved recall [60]. Large effects of understanding of the material to be learned on encoding efficiency is a well-established result in cognitive psychology [61], and this phenomenon is likely to at least partly explain the observed effect of visuo-constructive ability level on the developmental trajectory of extended delay visual recall.

### Effects of hippocampal structure

The major theories of long-term memory, such as the standard consolidation theory [62] and multiple trace theory [63], would predict that the efficiency of hippocampal processes affect consolidation over the time interval used in the extended delay recall conditions in present study. A structurally immature hippocampus would likely lead to less efficient re-activation of encoded memories and a less efficient hippocampal-cortical dialogue. In the framework of the multiple trace theory, this may be seen as less efficient establishment of multiple traces in the medial temporal lobe and the neocortex - and consequently steeper forgetting rates. The present finding of unique developmental effects for extended delay recall fit this hypothesis, further supported by the existence – although quite modest – of relationships with hippocampal sub-region structure. This interpretation fits with an earlier cross-sectional study with a sample overlapping the present one, where extended delay long-term visuo-constructive recall was related to hippocampal volume while performance over 30 minutes was related to prefrontal cortex thickness [2]. In adults, the relationship between hippocampal volume and memory performance is generally not robust [64], but may be stronger with longitudinal designs [65–67], in development [68] and with memory tests spanning longer time intervals [7]. While we could not find any developmental studies examining the relationship between hippocampal microstructure and memory in development, studies of adults suggest that MD may more closely than volume be associated with episodic memory performance [34, 37–39]. In the present study, four hippocampus-memory relationships were identified at a nominal α-level of .05, with two each for MD and volume. Applying a strict Bonferroni correction, not taking into account the correlations between the different variables, only the relationship between extended delay verbal recall and anterior MD survived. However, inspecting the heat map in Figure 4, it is difficult to make inferences about stronger relationships with memory for one of the structural hippocampal measures over the other.

Similarly, there were not obvious differences in the memory relationships for the hippocampal sub-regions. The role of aHC vs. pHC in episodic memory function depends on the specific [11–13] and general demands of the task, with aHC more involved in encoding and pHC in retrieval [11, 13–15]. Using only behavioral testing, we could not distinguish between encoding and retrieval effects on the extended delay memory scores. We have previously found that children engaged pHC more than aHC, while aHC was more activated relative to pHC already in teenagers [69]. The partly different structural developmental trajectories of the hippocampal sub-regions may have impact on more specific memory processes, which we were not able to detect with our behavioral task. Positive correlations between both aHC [24] and pHC [3] have been reported in development, as in adults [23, 25, 70]. The present results did not show consistent aHC-pHC differences. Rather, the effects varied as a function of retention interval, with tendencies for stronger effects for the extended delay recall tests than the 30 min tests, as discussed above.

### Limitations

There are multiple limitations of the current study. For the oldest participants, some approached the maximum score, which makes it possible that ceiling effects affected the results. Inspections of the scatterplots in Figure 2 do not indicate that this is a serious issue, but this may still have contributed to reduce the observed age-effects, especially among the oldest adolescents. Among the youngest participants, some showed low scores, which may suggest possible floor effects in this age-range. As the scores increased rapidly with advancing age, however, this has likely not affected the age-trajectories substantially. We did not directly correct for selective attrition or learning effects, which may impact the results [71–73]. Instead, we included subject time-point as a covariate in all analyses, which should effectively control for the effects of taking the test multiple times. Another difficult issue when studying long-term memory is how to deal with differences in initial learning rate. In the present study, scores on the 30 min condition were regressed out in the models including long-term memory, yielding good statistical control for differences is initial memory performance level. This is a challenging issue, however, with multiple possible solutions with different strengths and weaknesses. Some have matched initial performance levels by e.g. multiple repetitions of items during encoding. However, no consensus has been reached regarding whether or not degree of initial learning affects rate of forgetting, and there is presently no agreement about how best to tackle this problem (see [58] for an extensive discussion of these issues). Finally, the present study focused on the hippocampus. Obviously, brain structure - memory relationships could have been found in other brain regions not included in the present work.

## 5.0 Conclusion

Extended delay episodic memory recall showed unique developmental trajectories not accounted for by more immediate long-term memory. For visuo-constructive extended delay recall, these improvements during childhood and adolescence could be accounted for by general visuo-constructive ability level. This was not the case for verbal extended delay recall, where development was not affected by verbal ability. Finally, episodic memory over extended delays was modestly related to hippocampal structure. Experimental work, including functional brain activation studies, will be necessary to reveal to which degree development of episodic memories lasting days in contrast to hours or less are caused by maturation of consolidation processes versus encoding-related processes.

## Funding

This work was funded by the Department of Psychology, University of Oslo (to K.B.W., A.M.F.), the Norwegian Research Council (to K.B.W., A.M.F.) and the project has received funding from the European Research Council’s Starting/ Consolidator Grant schemes under grant agreements 283634, 725025 (to A.M.F.) and 313440 (to K.B.W.).

## Declarations of interest

Declarations of interest: none.

